# Bio-optical properties and radiative energy budgets in fed and starved scleractinian corals *(Pocillopora damicornis)* during thermal bleaching

**DOI:** 10.1101/377937

**Authors:** Niclas Heidelberg Lyndby, Jacob Boiesen Holm, Daniel Wangpraseurt, Christine Ferrier-Pagès, Michael Kühl

## Abstract

Corals achieve outstanding photosynthetic quantum efficiencies approaching theoretical limits (i.e. 0.125 O_2_ photon^-1^) and it is unknown how such photosynthetic efficiency varies with environmental stress. In this study, we investigated the combined effects of thermal stress and active feeding on the radiative energy budget and photosynthetic efficiency of the symbiont-bearing coral *Pocillopora damicornis* by using fiber-optic and electrochemical microsensors in combination with variable chlorophyll fluorescence imaging. At normal temperature (25°C), the percentage of absorbed light energy used for photosynthesis was higher for fed (~5-6% under low light exposure) compared to unfed corals (4%). Corals from both feeding treatments responded equally to stress from high light exposure (2400 μmol photons m^-2^ s^-1^), exhibiting a decrease in photosynthetic energy efficiency down to 0.5-0.6%. Fed corals showed increased resilience against thermal bleaching compared to unfed corals, as fed corals were able to uphold their high photosynthetic energy efficiency for 5 days longer during thermal stress, as compared to unfed corals, which decreased their photosynthetic energy efficiency almost immediately when exposed to thermal stress. We conclude that active feeding is beneficial to corals by prolonging coral health and resilience during thermal stress as a result of an overall healthier symbiont population.

## Introduction

Tropical coral reefs are among the most diverse and productive ecosystems on Earth. Reef-building corals thrive in an oligotrophic environment, where their success is largely driven by the endosymbiotic relationship with various clades of dinoflagellate microalgae in the genus *Symbiodinium*, collectively known as zooxanthellae, which are harbored inside endodermal host cells. The coral holobiont, i.e., the association of the coral cnidarian animal, *Symbiodinium*, and microbes associated with coral tissue and skeleton (Bourne et al., 2009), has an advantage in oligotrophic environments, as it is able to acquire heterotrophically fixed carbon and nutrients via prey capture and particle feeding, as well as gaining access to autotrophically fixed carbon via symbiont photosynthesis and subsequent leakage of photosynthates from the zooxanthellae to the host. While zooxanthellae only represent ~5-15 % of the coral tissue biomass (Odum & Odum, 1955; Thornhill *et al*., 2011), they provide up to 95% of the coral’s energy demand (Muscatine *et al*., 1981; Edmunds & Davies, 1986). However, corals are susceptible to environmental stress (e.g. changes in seawater temperature, nutrient stress, pH, O_2_, etc.), which can lead to the break-down of the coral-algal symbiosis and the expulsion of the symbiotic algae, a phenomenon known as coral bleaching (Lesser, 1996; Hoegh-Guldberg, 1999; Hoegh-Guldberg *et al*., 2007; Wiedenmann *et al*., 2012). Stress-induced mass bleaching events have increased in frequency due to global climate change, and such events now represent one of the greatest threats to coral reefs worldwide (Hoegh-Guldberg, 1999; Hughes *et al*., 2003, 2017; Ainsworth *et al*., 2016). On a larger scale, the most widely acknowledged stressors are instances of above-average sea water temperatures, due to global warming, in combination with excess solar irradiance (Glynn, 1996). The primary biochemical causes of coral bleaching are still debated (Brown & Dunne, 2015) but are strongly linked to the formation of reactive oxygen and nitrogen species in corals upon environmental stress (Lesser, 1996; Bou-Abdallah *et al*., 2006). Irradiance exposure plays a central role in the bleaching response of corals, and excess irradiance can e.g. lead to photodamage of photosystem II and essential repair systems in the algal symbiont leading to the production of harmful O_2_ radicals (Lesser, 1996; Lesser & Farrell, 2004; Hill *et al*., 2011). A better understanding of how corals handle light is thus important for a mechanistic description of coral bleaching.

In reef environments, supersaturating light levels can lead to photoinhibtion, i.e., the decrease in photosynthetic quantum efficiencies due to increasing damage to the photosystems and cellular repair mechanisms. Fluctuations in irradiance exposure (ranging from seconds to hours) thus create a need for regulating and optimizing light harvesting and utilization of incident light energy on coral reefs (Anthony & Hoegh-Guldberg, 2003; Veal *et al*., 2010b). Photon absorption and the light microenvironment in corals are strongly modulated by the optical properties of the coral tissue and skeleton (Wangpraseurt *et al*., 2012, 2014a; Marcelino *et al*., 2013). The density and distribution of coral host pigments and symbionts in the tissue, as well as the scattering properties of both coral tissue and skeleton are important factors for modulating *Symbiodinium* light absorption and thus photosynthetic efficiency *in hospite* (Wangpraseurt *et al*., 2014b; Gittins *et al*., 2015).

Photons traveling through coral tissue can undergo several scattering events, leading to localized scalar irradiance enhancement in upper tissue layers (Wangpraseurt *et al*., 2014a, 2017b). While such enhancements in light exposure contribute to the high photosynthetic efficiency of corals under optimal irradiance exposure (Brodersen *et al*., 2014), the same light enhancing mechanisms can cause stress when corals are subject to excess irradiance, eventually resulting in coral bleaching. Enhanced skeletal scattering from bleaching leads to enhanced light absorption by the remaining *Symbiodinium* cells and can thus induce further light stress, ultimately accelerating the bleaching response; a process known as the optical feedback loop (Enríquez *et al*., 2005; Wangpraseurt *et al*., 2017b).

Furthermore, the absorption of light energy by symbiont photopigments is a major driver of radiative heat generation in coral tissue (Jimenez *et al*., 2012), with the rate of coral heating being directly proportional to the amount of incident light energy being absorbed (Jimenez *et al*., 2008; Welch & van Gemert, 2011). Excess heat from the tissue is dissipated via convection into the surrounding water across a thermal boundary layer (TBL), and via heat conduction down into the coral skeleton (Jimenez *et al*., 2008). The thickness of the TBL changes depending on the ambient flow regime, coral topography, and shape (analogous to the well-known diffusive boundary layer, DBL), while coral tissue surface area/volume ratio affects coral surface warming and cooling (Jimenez *et al*., 2008, 2011). Coral surface topography thus affects the properties of the flow dependent DBL and TBL, ultimately controlling the exchange of solutes and heat (Lesser *et al*., 1994; Kaandorp *et al*., 1996; Jimenez *et al*., 2008; Chan *et al*., 2016).

The photosynthetic quantum efficiency of corals (in units of mol O_2_ produced or mol C fixed via photosynthesis per mol photons absorbed) describes how effectively incident irradiance absorbed by symbiont photopigments is used for photosynthesis and carbon fixation. The photosynthetic quantum efficiency is affected by environmental parameters such as irradiance, temperature, nutrient status and flow (Falkowski & Dubinsky, 1981; Roth & Deheyn, 2013; Brodersen *et al*., 2014). Corals respond to such environmental factors by modulating e.g. their morphology, symbiont density and pigment composition, as well as their feeding behavior depending on the availability of nutrients (Enríquez *et al*., 2005; Ferrier-Pagès *et al*., 2010; Jimenez *et al*., 2011). Phosphate starvation can e.g. alter *Symbiodinium* population density in corals, as an increase in the N:P ratio (e.g., by an increased dissolved inorganic nitrogen, DIN, supply) will increase symbiont density, while changing the lipid composition of algal membranes, leading to reduced photosynthetic efficiency and photodamage, especially at excess temperatures (Wiedenmann *et al*., 2012). Consequently, starved corals under thermal stress can be more prone to photoinhibition and bleaching, resulting in reduced photosynthesis and reduced chlorophyll content in coral tissues (Ferrier-Pagès *et al*., 2010).

Studies of photosynthetic efficiency and radiative energy budgets accounting for the fate of incident and absorbed light energy in benthic marine systems have so far focused on sediments, biofilms and microbial mats (Al-Najjar *et al*., 2010, 2012; Lichtenberg *et al*., 2017), with the exception of a single study on a symbiont-bearing coral (Brodersen *et al*., 2014). It was found that, while the majority (>96%) of incident light energy was absorbed and dissipated as heat, the quantum efficiency (Q_E_) of corals measured under low to moderate irradiance was close to the theoretical maximum of 0.125 mol O_2_ per mol photons, showing that light is used very efficiently for photosynthesis in corals (Brodersen *et al*., 2014). Furthermore, the proportion of photosynthesis decreased with increasing irradiance, while heat dissipation increased and reflectance remained unchanged at about 10% of the incident irradiance. Similar observations have been made in microbial mats, although these systems exhibit a much lower photosynthetic efficiency (Al-Najjar *et al*., 2010). A direct comparison of how the radiative energy budget and thus photosynthetic efficiency changes during bleaching in starved versus fed corals, has so far not been reported.

In this study and in an accompanying paper (Lyndby et al., submitted), we investigated the radiative energy and carbon budgets of starved and fed corals of *Pocillopora damicornis* during thermal bleaching. The present study focusses on the photosynthetic efficiency of *P. damicornis*, where we present closed radiative energy budgets for fed and starved corals during thermal stress-induced bleaching based on microsensor measurements, and variable chlorophyll fluorescence imaging. In an accompanying paper (Lyndby et al., submitted), we focus on the carbon budget of fed and unfed *P. damicornis* during their bleaching response.

## Methods

### Corals

*Pocillopora damicornis* corals were supplied by the Centre Scientifique de Monaco (CSM). Corals were prepared by cutting eight mother colonies into ~120 fragments of 2-3 cm in diameter, and hung from nylon threads in the tank. All corals were kept under a 12:12 hours day-night cycle using white light (250W metal halide lamps; Philips, Netherlands), at a downwelling photon irradiance (400-700 nm) of 200-250 μmol photons m^-2^ s^-1^, as determined by a quantum irradiance sensor connected to a light meter (Li-COR 250a, Li-COR, USA).

### Thermal stress and feeding treatments

Corals were exposed to 4 different experimental treatments: 1) fed and heated, 2) fed and not heated, 3) unfed and heated, 4) unfed and not heated. Eight tanks (water renewal rate of 10 L h^-1^, 25°C, salinity of 35) were set up two months in advance of the bleaching experiment as to divide the corals into two groups of 4 tanks, each with either fed or unfed corals. Fed corals were fed once a day four times per week with ~4000 *Artemia* nauplii per coral fragment, while unfed corals were kept unfed throughout the study, including the two months prior to measurements.

About 20 fragments were kept in each control tank, while three thermal stress tanks contained a total of 40 fragments for each feeding treatment. At the beginning of the study, all 8 tanks started at 25°C. The tanks used for thermal stress were then gradually ramped up to 30°C over a period of five days, increasing the temperature by 1°C per day. Thermally stressed corals were kept at 30°C for an additional two to three days before starting measurements. First measurements were performed on corals after the first two to three days of 30°C thermal stress (T_1_). Measurements were then performed after an additional five days of thermal stress (T_2_). Measurements on control corals (T_0_) were done in between the waiting period for T_1_ and T_2_.

### Experimental setup and approach

Microsensor measurements were performed with corals placed in a black acrylic flow chamber. The flow chamber was supplied with seawater (25°C, salinity 35) from a heated water reservoir (10 L) at a flow rate of ~0.25 cm s^-1^. A motorized micromanipulator (MU-1, PyroScience, Germany) was attached to a heavy-duty stand to facilitate positioning of microsensors on fragments in a 45° angle relative to the vertically incident light. A digital microscope (Dino-Lite Edge AM7515MZTL, AnMo Electronics Corporation, Taiwan) was used to carefully position the sensor tip on the coral tissue surface (Figure 1e-f). Corals were illuminated from above with white light provided by a tungsten halogen lamp (KL-2500 LCD, Schott, Germany), fitted with a fiber light guide and collimating lens (Supplementary figure 1). A calibrated spectroradiometer (MSC-15, GigaHertz-Optik, Germany) was used to quantify the absolute downwelling photon irradiance at different lamp settings (80, 167, 250, 480, 970, and 2400 μmol photons m^-2^ s^-1^), and in addition to record downwelling irradiance spectra in units of W m^-2^ nm^-1^. The irradiance was adjusted without changing spectral composition by adjusting a metal disk with varying perforation between the halogen light bulb and the fiber-optic light guide in the lamp light path. Setups were covered with black cloth during measurements. For each coral replicate used in this study, microsensor measurements were done on three randomly chosen polyps located on the branch tips (Figure 1).

**Figure 1.**
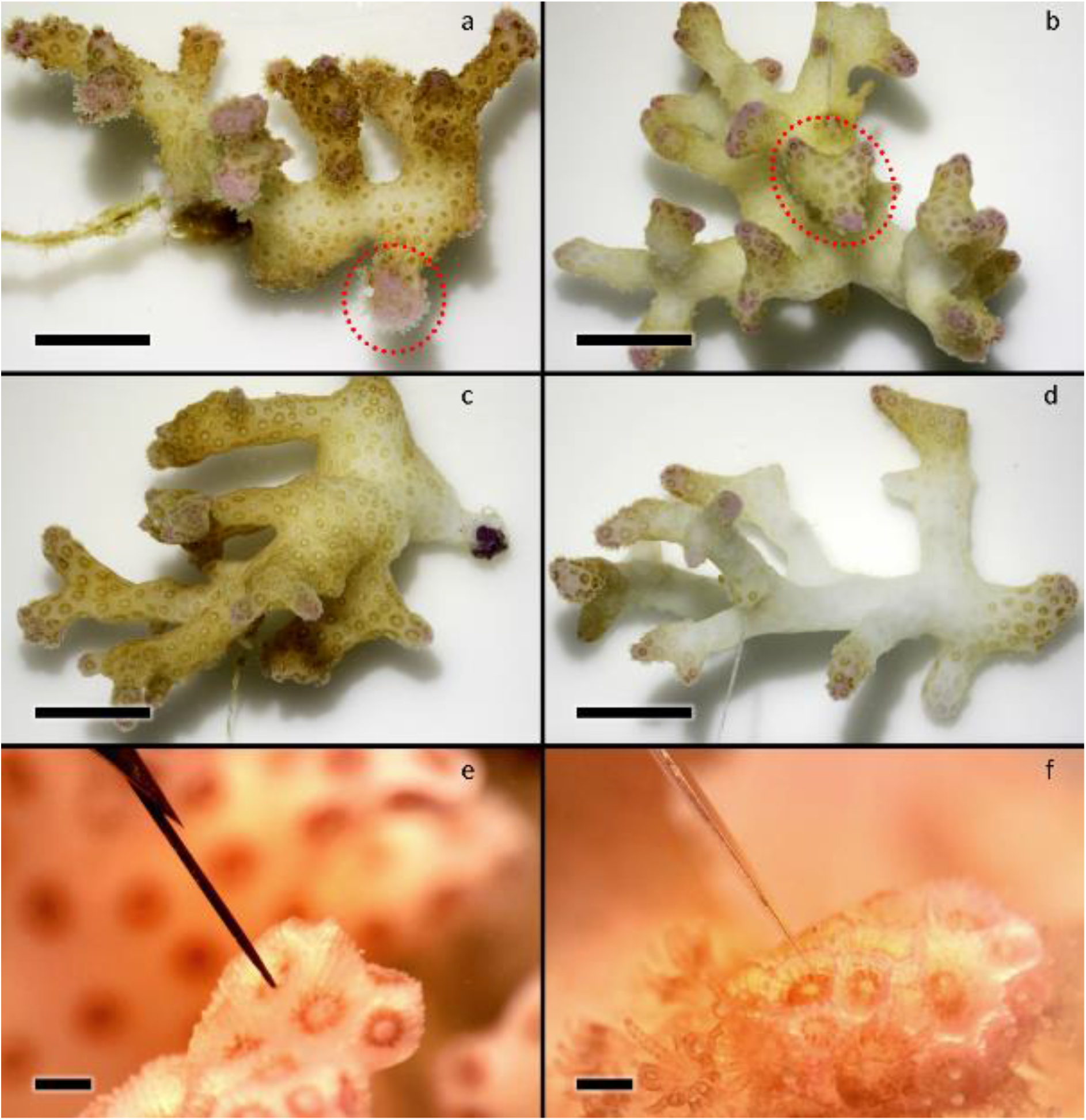
Representative photographs of *Pocillopora damicornis* corals subject to four experimental treatments: controls (a-b) and thermally stressed (c-d) fragments of fed (a, c) and unfed (b, d) *P. damicornis*. Photos were taken at the end of the experimental period, on the last day of measurements on unfed fragments at T_1_. All photos were normalized against a white standard by adjusting the brightness/contrast levels using ImageJ. Note that the coral animal tissue structure was intact for all fragments. The scale bar in panels a-d equals 1 cm in length, scale bar in panel e-f equals 2.5 mm in length. Red dotted circles on panels a-b indicate examples of areas used for microsensor measurements. Panels e-f show close-ups of measuring areas with scalar irradiance (e) and O_2_ (f) microsensors positioned on polyp tissue. The sensor tips were aimed at the highly pigmented polyp rims located at the branch tips of fragments.

### Microsensor measurements

#### Scalar Irradiance and reflectance measurements

Spectral scalar irradiance was measured with a fiber-optic scalar irradiance microsensor (spherical tip diameter ~50 μm; Rickelt *et al*., 2016). Measurements were performed at the tissue surface and within the coral tissue for corals with tissue thicker than 50 μm. Tissue thick enough (>50 μm) was carefully punctured with a tungsten needle (tip diameter ~1 μm, see Wangpraseurt *et al*., 2017b for details). after which the sensor tip was quickly moved into the tissue. A reference spectrum of the incident downwelling irradiance was measured afterwards without the coral fragment and with the scalar irradiance microsensor tip placed over a black light well at the same position in the flow chamber and incident collimated light field as the coral surface.

Irradiance reflectance of the coral tissue surface was measured with a 0.7 mm wide flat-cut fiber-optic reflectance probe. Coral reflectivity was measured with the probe positioned at a distance of 500 μm from the tissue surface. All reflectivity measurements were normalized with the reflectivity of a 99% white diffusing reflectance standard (Spectralon, Labsphere, USA) measured under identical configuration as on the coral surface, but performed in air with the same distance from the tungsten halogen lamp (KL 2500, Schott, Germany).

Both scalar irradiance and reflectance microprobes were connected to a fiber-optic spectrometer (USB 2000+, Ocean Optics, USA), and all spectra were recorded with the software SpectraSuite (version 2.0.162, Ocean Optics, USA) running on a PC connected to the spectrometer via an USB cable.

#### Photosynthesis and O_2_ measurements

Gross photosynthesis was measured with Clark-type O_2_ microelectrodes with a tip diameter of ~25 μm, a low stirring sensitivity (<5%), and a fast response time (<0.5 s; OX-25 fast, Unisense, Denmark). The microelectrodes were connected to a pA meter (Unisense, Denmark), and were linearly calibrated from readings in 100% air saturated seawater and anoxic water (by addition of Na_2_SO_3_) at experimental temperature and salinity. The corresponding O_2_ concentration (μmol L^-1^) in seawater at 100% air saturation was taken from gas tables for the O_2_ solubility in air-saturated water as a function of temperature and salinity (www.unisense.dk). Data were recorded on a strip-chart recorder (BD25, Kipp&Zonen, Netherlands) connected to the pA meter. Gross photosynthesis was estimated using the light-dark shift method as described in detail by Revsbech & Jørgensen (1983), where the measurements performed at the coral tissue surface were regarded as representative of the entire tissue volume of *P. damicornis* given that the tissue thickness in most places was ~100-200 μm thick (see results), which approximates the spatial resolution of the light-dark shift method during the 2-3 second period of darkening. Accordingly, the areal rate of gross photosynthesis (GPP) was estimated by multiplying the measured volume-specific gross photosynthesis at the tissue surface (in nmol O_2_ cm^-3^ s^-1^) by the local tissue thickness (in cm). The local tissue thickness was estimated from the difference in depth position of the scalar irradiance measurements done at the water-tissue and tissue-skeleton interface, respectively.

#### Temperature measurements

Coral surface tissue heating was measured with a temperature microsensor (tip diameter ~50 μm; TP50, Unisense, Denmark) connected to a thermocouple meter (T301, Unisense, Denmark). The thermocouple meter was interfaced to a PC via an A/D converter (DCR 16, PyroScience, Germany) connected to a PC running the software Profix (version 4.51, PyroScience, Germany) that controlled the positioning of the micromanipulator and data acquisition. The temperature microsensor was linearly calibrated against a high precision thermometer (Testo 110, Testo AG, Germany) in seawater at 19°C and 25°C. Temperature profiles were measured from the tissue surface into the ambient water in steps of 100-300 μm (with a resting time of 7 seconds at each depth position) across the TBL (Jimenez *et al*., 2008). The thickness of the TBL was determined as the intersection of the linear part of the temperature profile and the ambient water (see figure 1b in Jimenez *et al*., 2008). ΔT was determined as the temperature difference in °C between the coral surface and ambient water outside the TBL (i.e., 5000 μm from the tissue surface). All temperature measurements were done under a high incident photon irradiance of 2400 μmol photons m^-2^ s^-1^.

### Chlorophyll content and symbiont density

Chlorophyll *a*+*c*_2_ content and symbiont density were determined for 4-5 coral fragments from each treatment at each time point. For this, tissue was first detached using a Water Pick with filtered seawater (FSW, 0.45 μm pore size). The tissue slurry was then homogenized using a Potter tissue grinder. For chlorophyll content determination, 5 mL of the homogenate was centrifuged at 11000g for 15 minutes at 4°C, where after the supernatant was discarded and the remaining algal pellet was re-suspended in 5 mL pure acetone. Pigments were extracted at 4°C over a period of 24 h in darkness, before they were centrifuged. The pigment-containing supernatant was collected and the chlorophyll content was determined by the spectrophotometric method of Jeffrey & Humphrey (1975) using a spectrophotometer (UVmc2, Safas, Monaco).

The density of zooxanthellae was determined by centrifugation of 0.1 mL of the tissue homogenate at 850g for 10 minutes. The supernatant was discarded, and the algal pellet was re-suspended in FSW that was subsequently used for ten separate chamber counts. The zooxanthellae were counted according to the method described by Rodolfo-Metalpa et al. (2006) using image analysis software (Histolab 5.2.3, Microvision Instruments, France).

Values of chlorophyll content and symbiont density were normalized against the skeleton surface area of the individual coral fragments as determined by the wax dipping method described by Veal *et al*. (2010a).

### Imaging of variable chlorophyll fluorescence and absorptivity

The absorptivity and PSII quantum yield of randomly selected fragments from both control tanks T_0_, unfed T_1_, and fed T_2_ were measured using a variable chlorophyll fluorescence imaging system (I-PAM, IMAG-MIN/R, Walz, Germany; Ralph *et al*. (2005). The measuring light intensity was adjusted for each measurement to yield a F_0_ value of about 0.1 for each sample. Data was collected by placing 3-4 fragments from one treatment and time point in a clear glass container filled with seawater from the holding tank. Fragments were dark acclimated for 5 minutes before measuring the maximal PSII quantum yield (F_v_/F_m_) by applying a strong saturation pulse (>3000 μmol photons m^-2^ s^-1^ for 0.8 s) with the imaging PAM, calculated as:

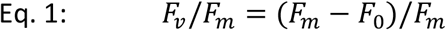

Rapid light curves (RLC) were performed for actinic light intensities ranging between 0-1600 μmol photons m^-2^ s^-1^. For each RLC, corals were exposed to a total of 14 light intensities (12, 40, 73, 99, 132, 162, 190, 292, 437, 606, 853, 1177, 1631 μmol photons m^-2^ s^-1^) for 10 seconds at each light intensity. The effective PSII quantum yield, Y(II), quantum yield of regulated energy dissipation, Y(NPQ), and quantum yield of nonregulated energy dissipation, Y(NO), were calculated as:

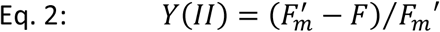

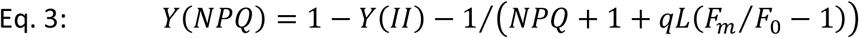

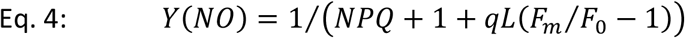

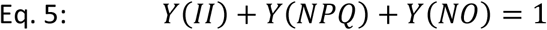

See Hill *et al*. (2004) and Baker (2008) for more details on variable chlorophyll fluorimetry. Calculations were performed for defined tissue areas by selecting regions of interest using the software ImagingWin (v2.41a, Walz, Germany). See Table 1 for definitions of parameters used.

**Table 1.** Definitions of parameters used for imaging of variable chlorophyll fluorescence.

| Parameter | Definition |
| --- | --- |
| $F_m$ | Maximal fluorescence yield (dark adapted) |
| $F_0$ | Minimal fluorescence yield (dark adapted) |
| $F_v/F_m$ | Maximal PSII quantum yield |
| $F_m'$ | Maximum fluorescence yield (light adapted) |
| $F$ | Fluorescence yield (light adapted) |
| $qL$ | Fraction of PSII centers that are open |
| $Y(II)$ | Effective PSII quantum yield |
| $Y(NPQ)$ | Quantum yield of regulated energy dissipation |
| $Y(NO)$ | Quantum yield of nonregulated energy dissipation |

### Radiative energy budget calculations

The radiative energy budget was calculated following procedures described in detail by Al-Najjar *et al*. (2010) but with minor changes as described below. The absolute downwelling spectral irradiance (in W m^-2^ nm^-1^) was measured with a calibrated spectroradiometer (MSC15, GigaHertz-Optik, Germany), and integrated over 400-700 nm (PAR) to quantify the total incident radiant energy flux (J_IN_, J m^-2^ s^-1^).

From J_IN_, we estimated the proportion of light energy absorbed (J_abs_) and reflected (R) by the coral. The PAR irradiance reflectance (R_(PAR)_) was calculated as:

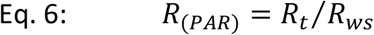

where *R_t_* is the coral tissue reflectivity, and *R_ws_* is the reflectivity measured from a 99% white diffusing standard (Spectralon, Labsphere, USA). The upwelling reflected light energy (R; in J m^-2^ s^-1^) was calculated as:

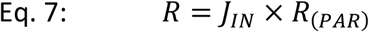

The light energy absorbed by the coral tissue, J_abs_ (in J m^-2^ s^-1^), was then calculated as the vector irradiance:

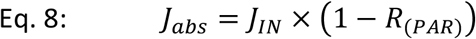

The absorbed light energy in the coral is either dissipated as heat, J_H_, or conserved by photosynthesis, J_PS_. The energy conserved by photosynthesis was estimated by recalculating the measured areal gross photosynthesis, GPP, in energy units by multiplication with the Gibbs free energy, *E_G_* = 482.9 *kJ* (*mol O*_2_)^-1^, i.e., the amount of energy produced during oxygenic photosynthesis (Al-Najjar *et al*., 2010). The total amount of energy conserved by photosynthesis, J_PS_ (in J m^-2^ s^-1^) was thus calculated as:

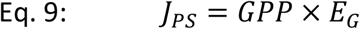

The remaining part of the absorbed light was dissipated as heat, J_H_, via an upward heat flux across the TBL or via a downward heat flux into the coral skeleton (Jimenez *et al*., 2008). The upward heat flux, J_H-up_, was calculated from temperature microsensor measurements across the TBL, using the linear slope of the temperature profile (K m^-1^) over the coral tissue surface, and multiplying it with the thermal conductivity of seawater at a salinity of 35, *k* = 0.6 *W m*^-1^ *K*^-1^:

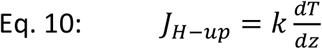

where *T* is the temperature measured in Kelvin, and *z* is the distance measured in m. It was not possible to directly measure the heat conduction from tissue into the coral skeleton with the fragile temperature microsensors, and the downward heat flux, J_H-down_ (in J m^-2^ s^-1^) was thus estimated as:

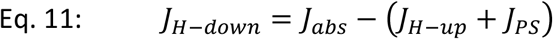

The total amount of energy dissipated as heat in the coral, J_H_ (in J m^-2^ s^-1^), was then calculated as:

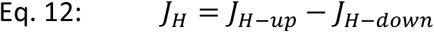

The entire radiative energy budget (in J m^-2^ s^-1^) is thus found as

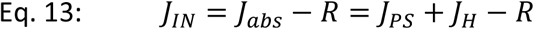

Energy budgets at lower irradiance irradiances were calculated based on detailed gross photosynthesis measurements on fragments from all time points (T_0_-T_2_) at previously mentioned photon irradiances (see Experimental setup and approach). The detailed measurements were combined with temperature and reflectance data, based on the assumption of a positive linear relationship between incident downwelling photon irradiance and heat dissipation (Jimenez *et al*., 2008), and a near constant percentage of reflection from the coral surface regardless of the incident irradiance (Al-Najjar *et al*., 2010; Brodersen *et al*., 2014). See Table 2 for definitions and units of abbreviations and parameters used.

**Table 2.** Abbreviations, definitions, and units used for the radiative energy budget calculations.

| Abbreviation | Definition | Unit |
| --- | --- | --- |
| ***PAR*** | Photosynthetically active radiation (400-700 nm) |  |
| ***TBL*** | Thermal boundary layer |  |
| ***R<sub>t</sub>*** | Tissue reflectivity | Arbitrary unit |
| ***R<sub>ws</sub>*** | White standard reflectivity | Arbitrary unit |
| ***R<sub>(PAR)</sub>*** | PAR irradiance reflectance ($R_t / R_{ws}$) | Arbitrary unit |
| ***J<sub>IN</sub>*** | Downwelling irradiance in energy units | J m<sup>-2</sup> s<sup>-1</sup> |
| ***R*** | Reflected light energy | J m<sup>-2</sup> s<sup>-1</sup> |
| ***J<sub>abs</sub>*** | Absorbed light energy | J m<sup>-2</sup> s<sup>-1</sup> |
| ***GPP*** | Areal rate of gross primary production | nmol O<sub>2</sub> cm<sup>-2</sup> s<sup>-1</sup> |
| ***E<sub>G</sub>*** | Gibbs energy | kJ (mol O<sub>2</sub>)<sup>-1</sup> |
| ***J<sub>PS</sub>*** | Energy conserved by photosynthesis | J m<sup>-2</sup> s<sup>-1</sup> |
| ***k*** | Thermal conductivity of seawater | W m<sup>-1</sup> K<sup>-1</sup> |
| ***dT/dz*** | Temperature gradient in the TBL | K μm<sup>-1</sup> |
| ***J<sub>H-up</sub>*** | Upward heat flux in energy units | J m<sup>-2</sup> s<sup>-1</sup> |
| ***J<sub>H-down</sub>*** | Downward heat flux in energy units | J m<sup>-2</sup> s<sup>-1</sup> |
| ***J<sub>H</sub>*** | Total heat flux in energy units | J m<sup>-2</sup> s<sup>-1</sup> |

## Results

### Variable chlorophyll fluorescence imaging

The effective PSII quantum yield, Y(II), at E_d_ = 1630 μmol photons m^-2^ s^-1^ in unfed corals did not change over time from (T_0_) and during thermal stress (T_1_), while fed corals showed a significant 3.2-fold increase of Y(II) after 8 days at 30°C (T_2_) relative to the control treatment (ANOVA, F(1, 46) = 12.7, P ≪ 0.01; Figure 2a). Overall, fed corals presented a higher Y(II) than unfed corals during all measurements (Figure 2a).

**Figure 2.**
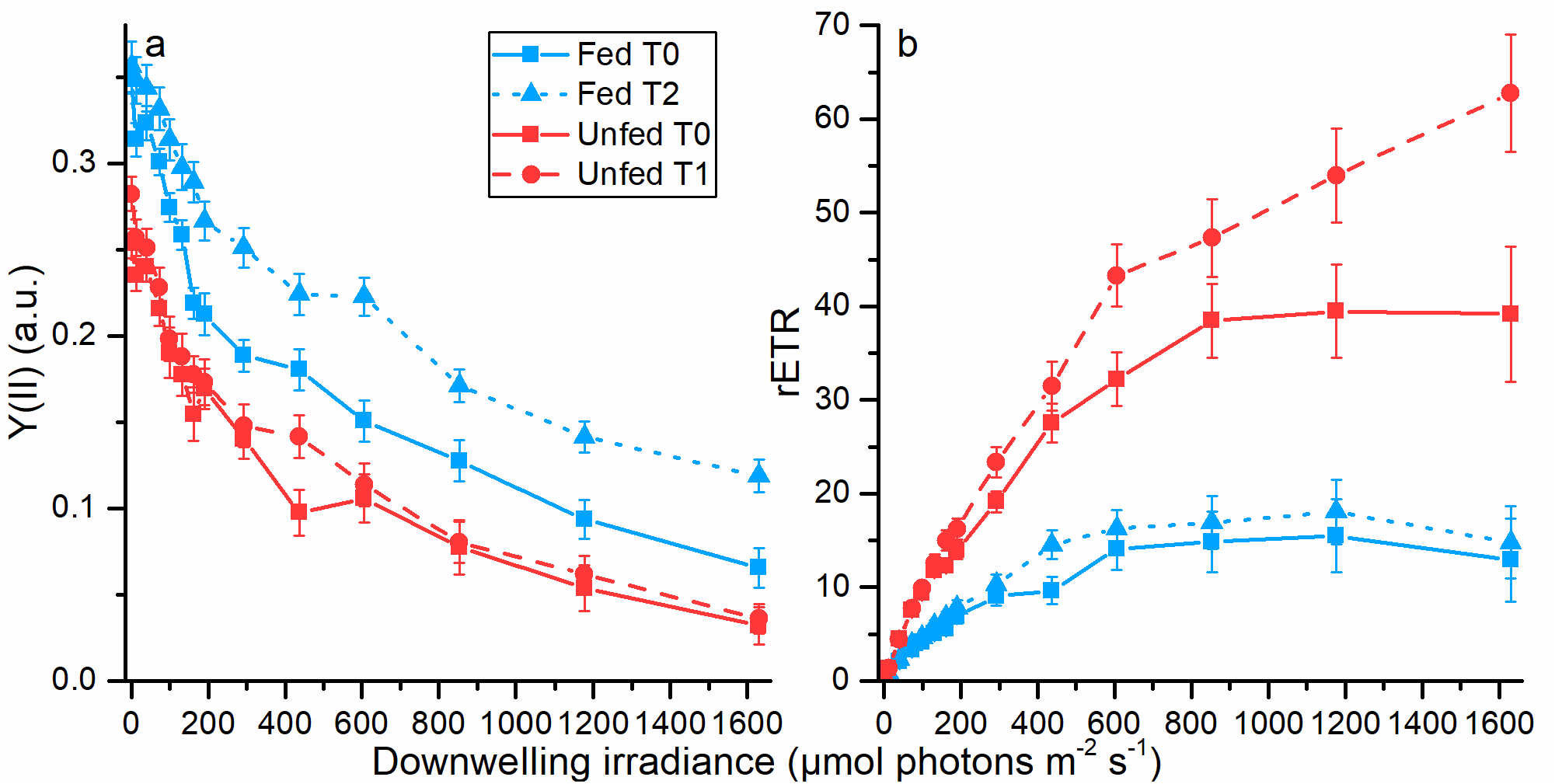
Variable chlorophyll fluorescence imaging of polyp tissue in fed and unfed *Pocillopora damicornis* during thermal stress treatment. (a) Effective quantum yield of PSII, and (b) Relative electron transport rate (rETR) before (T_0_) and after thermal stress (T_1_-T_2_). Symbols with error bars indicate means ± SE of different areas of interest in the images (n=24-36).

The maximal relative PS electron transport rate, rETR, at the highest photon irradiance (E_d_ = 1630 μmol photons m^-2^ s^-1^) did not differ between unfed corals before (T_0_) and after thermal stress (T_1_), while fed corals showed a significant 1.6-fold increase in rETR after 8 days at 30°C (T_2_) relative to the control (T_0_, ANOVA, F(1, 46) = 6.1, P = 0.02; Figure 2b). The rETR of fed corals was also 2-fold higher than the rETR of unfed corals, at both T_0_ and T_1_, and all light levels, which is in agreement with a higher F_v_/F_m_ (ANOVA, F(1, 106) = 44.3, P ≪ 0.01).

### Chlorophyll content and symbiont density

Zooxanthellae cell density at T_0_ was about 1.9 times higher for fed fragments compared to unfed fragments, and cell density decreased steadily in corals under both feeding treatments during thermal stress (T_1_-T_2_, Figure 3a). Relative to the starting population at T_0_, cell density loss in fed fragments was 10.1% after 3 days at 30°C (T_1_), and 27.5% after 8 days at 30°C (T_2_; ANOVA, F(2, 11) = 6.2, P = 0.016). For unfed fragments, cell density loss was 23.4% after 3 days at 30°C (T_1_), relative to the starting population (ANOVA, F(1, 7) = 9.1, P = 0.020; Figure 3a). The chlorophyll content was about 3.4 times higher in fed fragments as compared to unfed fragments at T_0_ (ANOVA, F(1, 21) = 17.8, P ≪ 0.01). No significant changes in chlorophyll content were observed during thermal stress in fragments from both feeding treatment relative to the starting content (T_0_-T_2_; ANOVA, F(2, 11) = 0.8, P = 0.49 for fed, F(1, 7) = 0.3, P = 0.61; Figure 3b).

**Figure 3.**
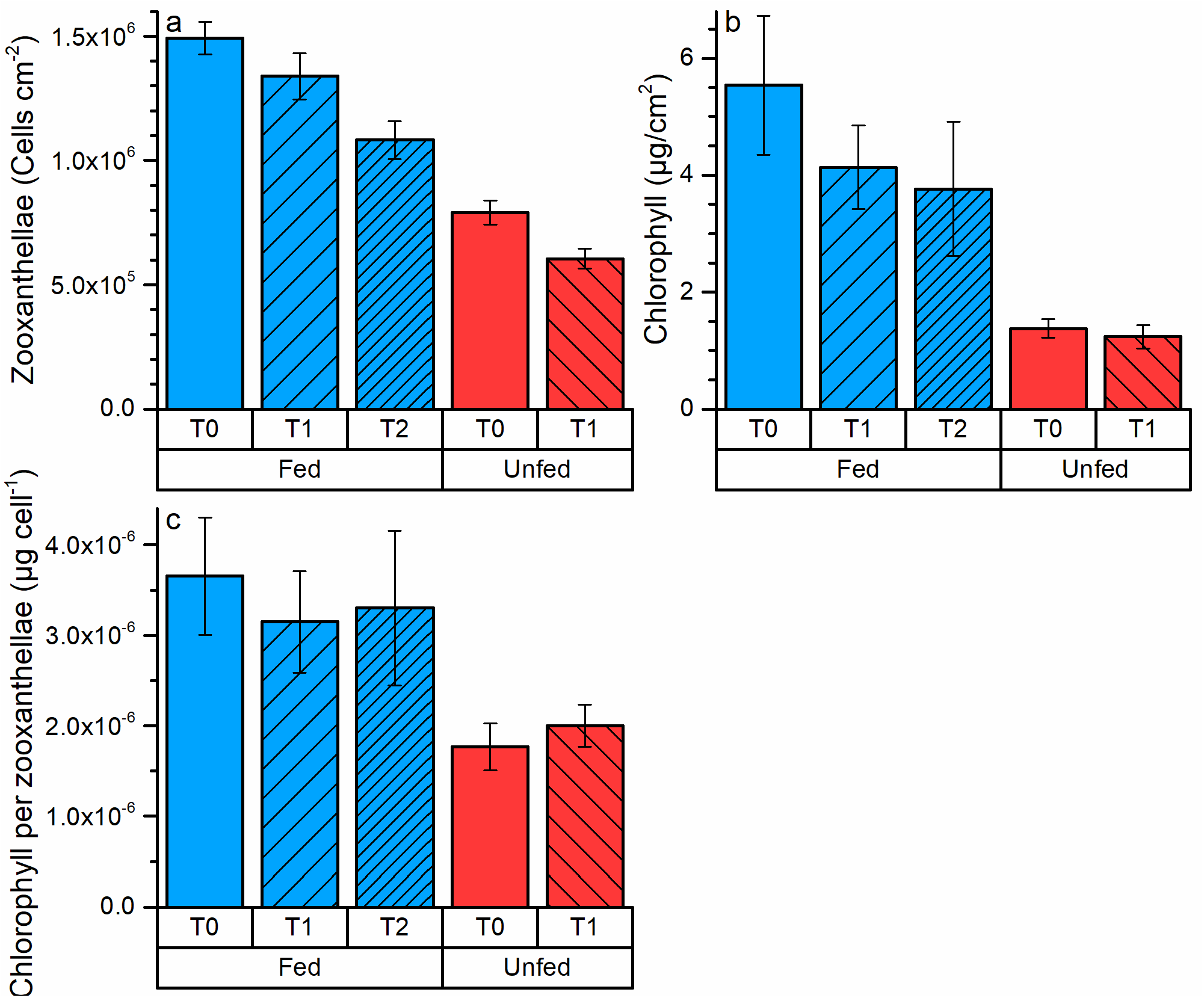
Mean zooxanthellae density (a), mean chlorophyll density (b), and ratio of total chlorophyll per zooxanthellae (c) in fed and unfed *Pocillopora damicornis* before (T_0_) and after thermal stress (T_1_-T_2_). Columns with error bars indicate means ± SE (n=4-5).

Chlorophyll content per cell was not affected by thermal stress relative to the starting content, but was on average 1.8 times higher in zooxanthellae from fed fragments compared to zooxanthellae from unfed fragments across all time points (T_0_-T_2_; ANOVA, F(1, 3) = 49.1, P < 0.01; Figure 3c).

### Scalar irradiance

Coral tissue surface photon scalar irradiance (425-700 nm) at T_0_ was 125.9% (±0.10 SE of mean) and 118.6% (±0.11 SE of mean) of the incident downwelling irradiance (E_d_ = 2400 μmol photons m^-2^ s^-1^) for fed and unfed corals, respectively (Figure 4a-b). Enhancement of tissue surface scalar irradiance peaked after 3 days at 30°C (T_1_) in both treatments (Figure 4a-b), reaching values of 140.9% (±0.12 SE of mean) and 146.7% (±0.07 SE of mean) relative to Ed for unfed and fed fragments, respectively. At T_0_, scalar irradiance at the coral tissue-skeleton interface was 95.3% (±0.08 SE of mean) and 110.6% (±0.08 SE of mean) of the incident downwelling irradiance for fed and unfed corals, respectively. Scalar irradiance at the tissue-skeleton interface did not change with thermal stress (T_1_-T_2_) relative to T_0_ (ANOVA, F(2, 34) = 1.5, P = 0.24 for fed, F(1, 21) = 2.8, P = 0.19 for unfed; Figure 4b).

**Figure 4.**
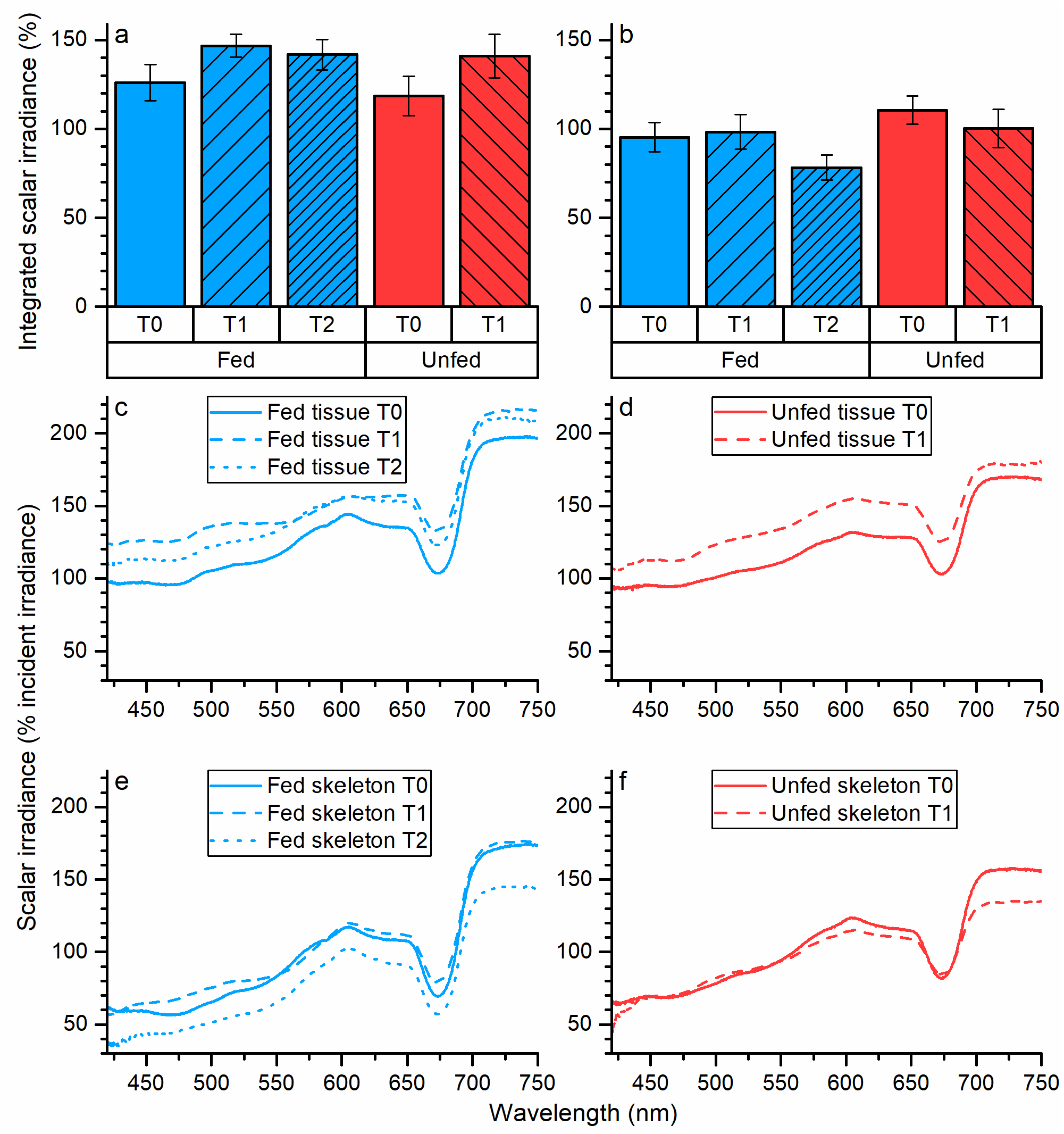
Scalar irradiance (a-b; integrated over 425-700 nm) in % of downwelling irradiance at the coral tissue surface (a) and the coral tissue-skeleton interface (b) of fed and unfed *Pocillopora damicornis* during thermal stress, and spectral scalar irradiance (c-f; in % of the incident downwelling irradiance) measured in polyp surface tissue (c-d) and at the coral skeleton-tissue interface (e-f) of fed (c, e) and unfed (d, f) fragments of *P. damicornis* during thermal stress. Columns with error bars and all spectra indicate means ± SE (n=10-15) for each time point per treatment (relative error of 20-50% for all spectra; c-f). Wavelengths below 425 nm were omitted due to high amounts of stray light in the spectrometer. Note that error bars in panel c-f are omitted for clarity, and y-axis start at 30%.

A distinct spectral attenuation was observed in the near infrared part of the spectrum (NIR, 700-750 nm) at tissue-skeleton interface in unfed corals after 3 days at 30°C (T_1_), and in fed corals after 8 days at 30°C (T_2_; Figure 4e-f) suggestive of the presence of endoliths in the coral skeleton, but no such endoliths were perceivable to the naked eye.

### Reflectance measurements

Spectral reflectance of PAR (400-700 nm) was 11.7% (±0.009 SE of mean) and 15.5% (±0.007 SE of mean) at T_0_ for fed and unfed fragments of *P. damicornis*, respectively (Figure 5a). Fed coral fragments showed a significantly increased reflectance at T_1_ and T_2_, relative to control measurements (T_0_; ANOVA, F(2, 45) = 3.5, P = 0.038), while unfed fragments showed no significant increase in reflectance between T_0_ and T_1_ (ANOVA, F(1, 34) = 0.6, P = 0.46; Figure 5a). Reflectance spectra from both fed and unfed treatment fragments showed lowest reflection in areas of absorption maxima for Chl *a* (435-440 and 675 nm), the PCP complex (540 nm), and Chl *c* (630-635 nm; Figure 5b-c).

**Figure 5.**
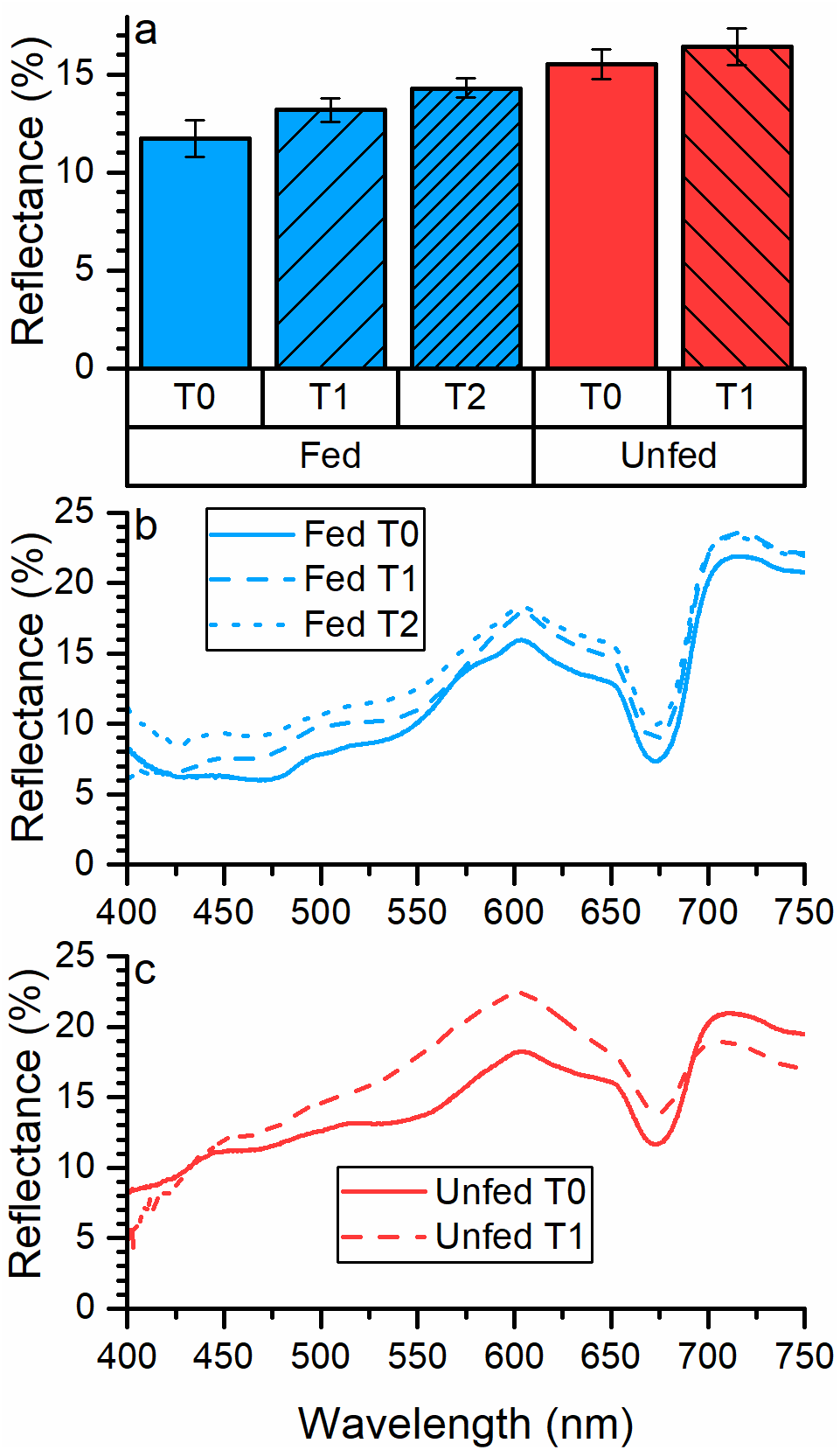
Diffuse reflectance (a; integrated between 400-700 nm), and spectral reflectance (b-c) of fed and unfed *Pocillopora damicornis* during thermal stress. Columns with error bars, and spectra indicate means ± SE (n=9-23). Note that error bars in panel b-c are omitted for clarity (SE <3%).

### Gross photosynthesis

Areal gross photosynthesis rates measured at high incident photon irradiance of Ed = 2400 μmol photons m^-2^ s^-1^ were not statistically different between measurements performed at any time point or feeding treatment (ANOVA, F(2, 19) = 1.4, P = 0.26 for fed, and F(1, 22) = 1.1, P = 0.30 for unfed; Figure 6a).

**Figure 6.**
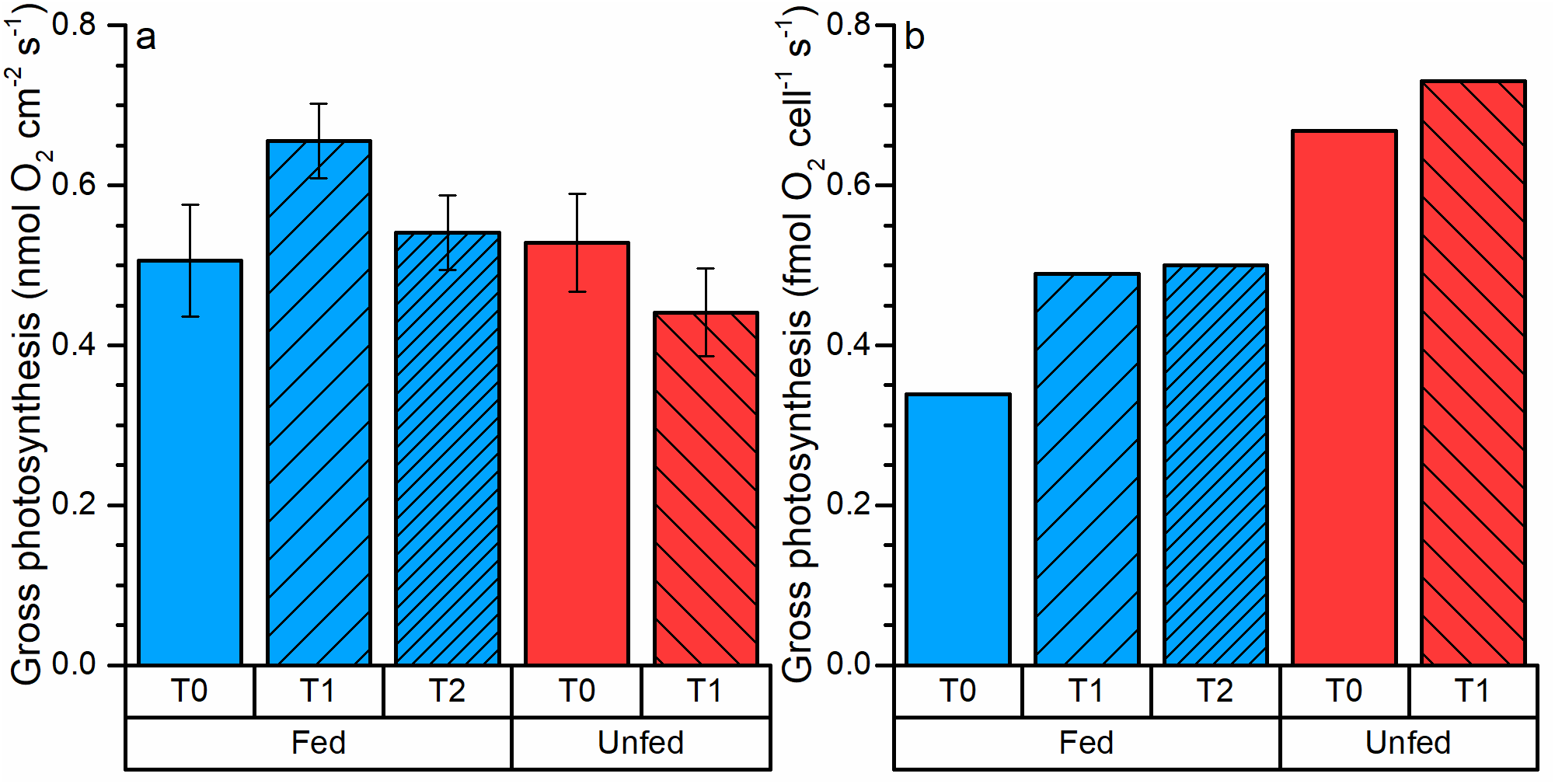
(a) Areal gross photosynthesis of fed and unfed specimens of *Pocillopora damicornis* under thermal stress measured at 2400 μmol photons m^-2^ s^-1^, averaged for each time point per treatment. Columns with error bars indicate means ± SE (n=5-12). (b) Cell-specific gross photosynthesis in fed and unfed *P. damicornis* for each treatment and time point.

For fed fragments, mean gross photosynthesis per cell was 1.4 times higher at T_1_ relative to control fragments at T_0_. No difference was observed for fed fragments between T_1_ and T_2_. For unfed fragments, gross photosynthesis per cell was 1.1-fold higher for T_1_ compared to T_0_ (Figure 6b). Unfed fragments had an overall higher gross photosynthesis per cell compared to fed fragments across all time points by ~0.25-0.3 times higher (Figure 6b).

Gross photosynthesis *versus* photon irradiance curves were corrected for the actual *in vivo* photon scalar irradiance measured with microsensors at the coral tissue surface for the individual treatments and time points (see methods for details). Areal gross photosynthesis for unfed control corals reached an asymptotic saturation level at P_max_ = 0.52 nmol O_2_ cm^-2^ s^-1^ (±0.01 SE, α = 0.0013±0.0006; T_0_) at a photon scalar irradiance of ~2150 μmol photons m^-2^ s^-1^ (Figure 7b). Fed control corals reached a similar saturation level at P_max_ = 0.53 nmol O_2_ cm^-2^ s^-1^ (±0.01 SE, α = 0.0020±0.0011; T_0_) at a photon scalar irradiance of ~2300 μmol photons m^-2^ s^-1^ (Figure 7a). P_max_ for unfed corals dropped to 0.44 nmol O_2_ cm^-2^ s^-1^ (±0.02 SE, α = 0.0007±0.0007; T_1_), while P_max_ for fed corals increased to 0.67 nmol O_2_ cm^-2^ s^-1^ (±0.03 SE, α = 0.0019±0.0017; T_1_) after 3 days of thermal stress, under a photon scalar irradiance of ~2600 μmol photons m^-2^ s^-1^ (Figure 7b). P_max_ decreased to 0.50 nmol O_2_ cm^-2^ s^-1^ (±0.03 SE, α = 0.0012±0.0014; T_2_) for fed corals after 8 days of thermal stress, and reached saturation at ~2200 μmol photons m^-2^ s^-1^ (Figure 7a).

**Figure 7.**
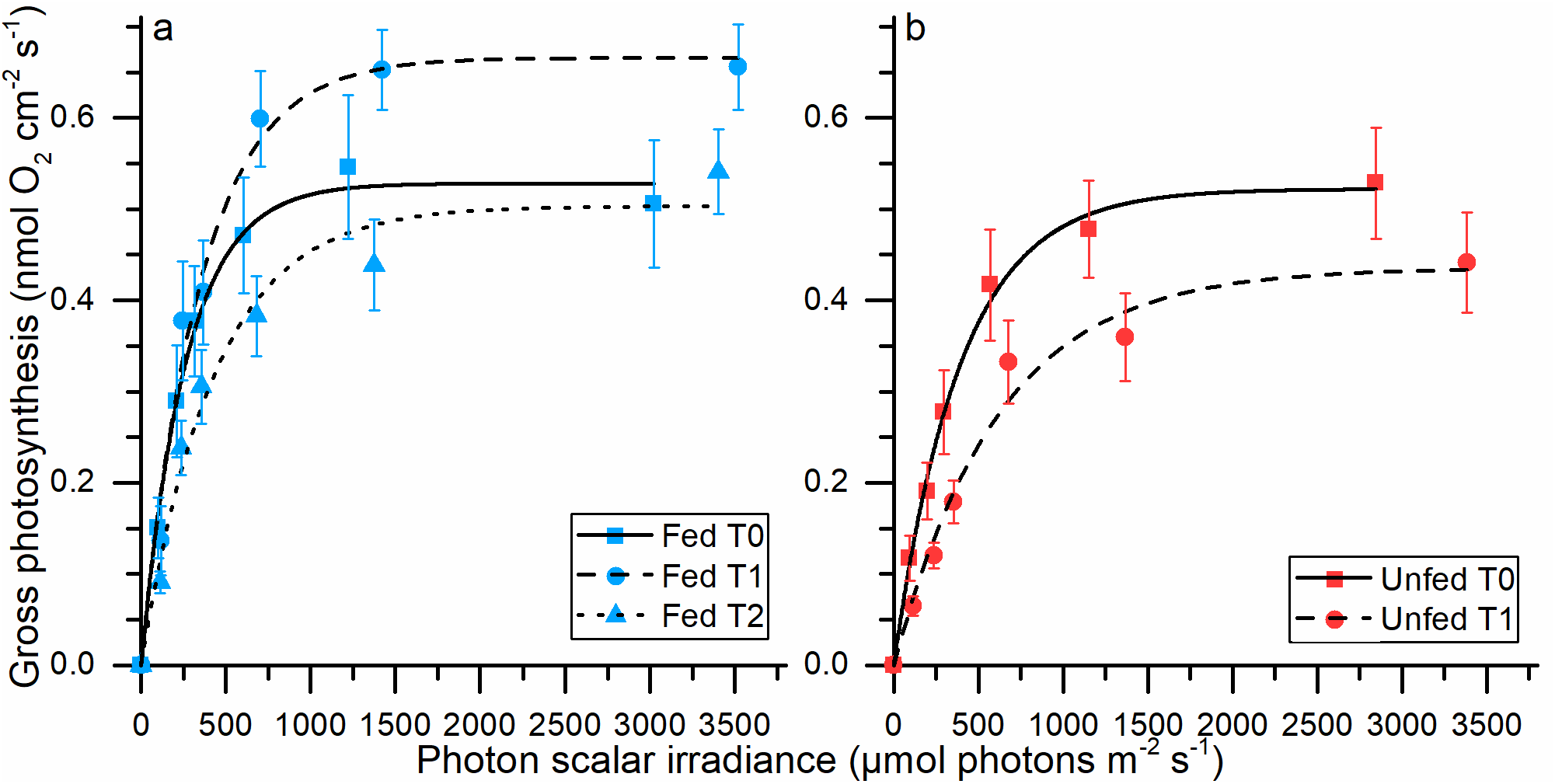
Gross photosynthesis *versus* photon scalar irradiance (PAR, 400-700 nm) curves for fed (a), and unfed (b) fragments of *Pocillopora damicornis* under thermal stress. Curves represent curve fits of an exponential model (Webb et al., 1974; R^2^>0.97-0.99). The light levels represent the actual amount of photons available for the individual time point and treatment by multiplying the downwelling irradiance with the local PAR enhancement obtained from the respective integrated scalar irradiance values (see Figure 4a). Error bars indicate ± SE of mean (n=5-12).

### Temperature measurements

The thermal boundary layer thickness of *P. damicornis* was 746 μm (±67 μm; SE of mean) for fed fragments and 617 μm (±37 μm; SE of mean) for unfed fragments under an incident photon irradiance of Ed = 2400 μmol photons m^-2^ s^-1^ and a flow velocity of ~0.25 cm s^-1^ (Figure 8a). For both fed and unfed fragments, surface heating at T_0_ reached a ΔT of 0.19°C (±0.02 SE of mean) and 0.17°C (±0.02 SE of mean) for fed and unfed corals, respectively (Figure 8b).

**Figure 8.**
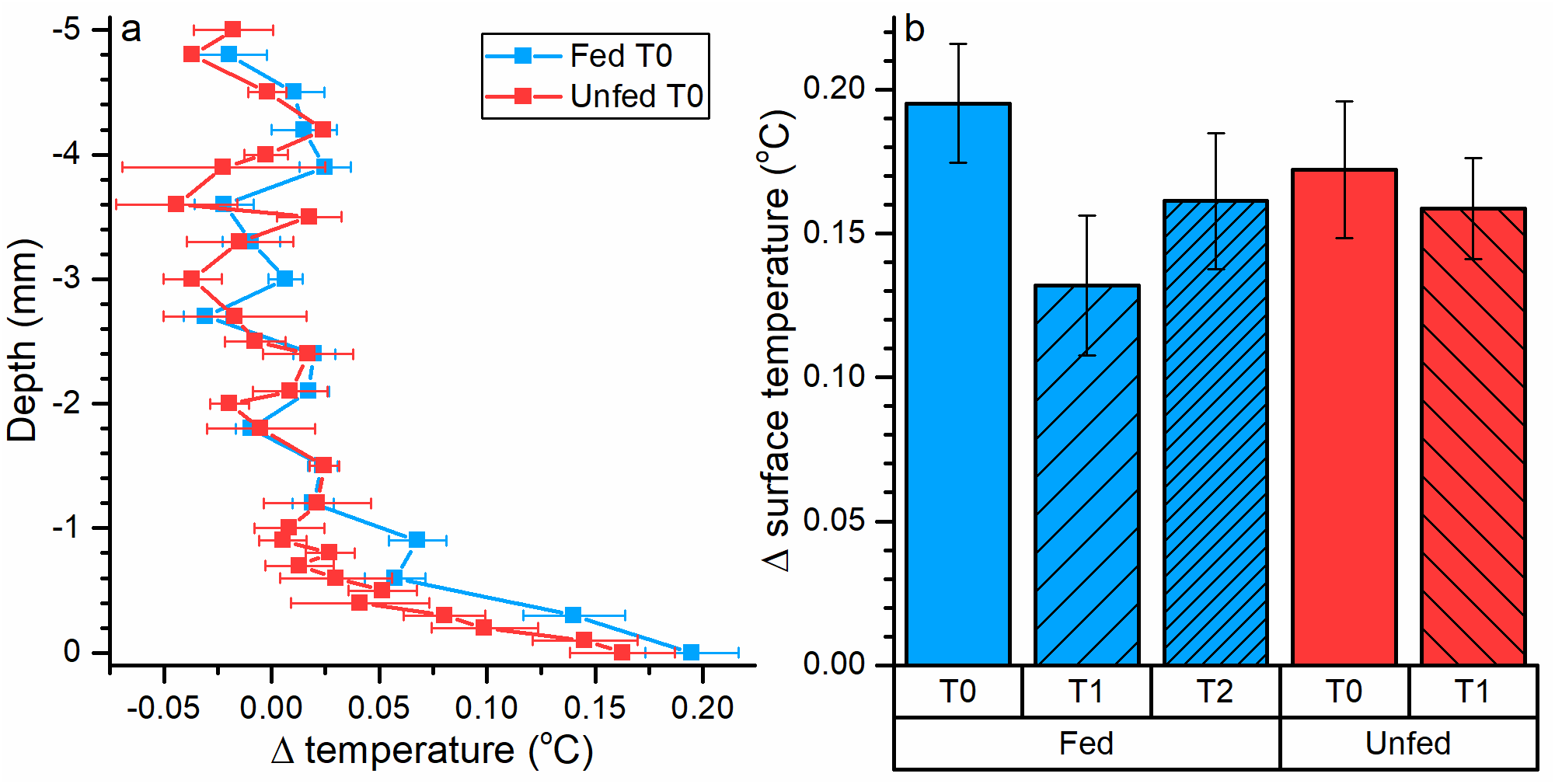
(a) Temperature profiles measured towards the tissue surface of fed and unfed control fragments of *Pocillopora damicornis* (T_0_, see T_1_-T_2_ in Supplementary figure 3). The x-axis shows the temperature difference between the coral tissue surface (0 mm) and the mean ambient water temperature under a downwelling photon irradiance of 2400 μmol photons m^-2^ s^-1^. Symbols with error bars indicate means ± SE of mean (n=9-15). (b) Mean temperature differences between the coral polyp tissue surface and the ambient water at 2400 μmol photons m^-2^ s^-1^. Columns with error bars indicate means ± SE (n=9-15).

### Radiative energy budget

We calculated radiative energy budgets based on microsensor measurements of reflection, gross photosynthesis, and temperature for an incident downwelling photon irradiance of 2400 mol photons m^-2^ s^-1^, which was equivalent to an incident irradiance (J_IN_) of 485.68 J m^-2^ s^-1^.

The amount of energy lost by tissue surface reflection increased with thermal stress, thus decreasing the amount of light absorbed by the coral tissue. For fed corals, reflected light energy was 11.72% (of the incident irradiance) at T_0_, 13.17% at T_1_, and 14.38% at T_2_. Likewise, reflectance increase in unfed corals from 15.01% of incident irradiance at T_0_, to 16.41% at T_1_ (Figure 9a). Under high irradiance, photosynthesis accounted for only 0.65% (±0.05 SE of mean; n = 3) and 0.57% (±0.05 SE of mean, n = 2) of the absorbed light energy in fed and unfed corals, respectively. We found no major differences in the amount of light energy conserved by photosynthesis, J_PS_, before and after thermal stress in either fed or unfed fragments (Figure 9b), and heat dissipation, J_H_, thus accounted for >99% of the absorbed energy dissipation in both fed and unfed fragments under high irradiance (Figure 9b).

**Figure 9.**
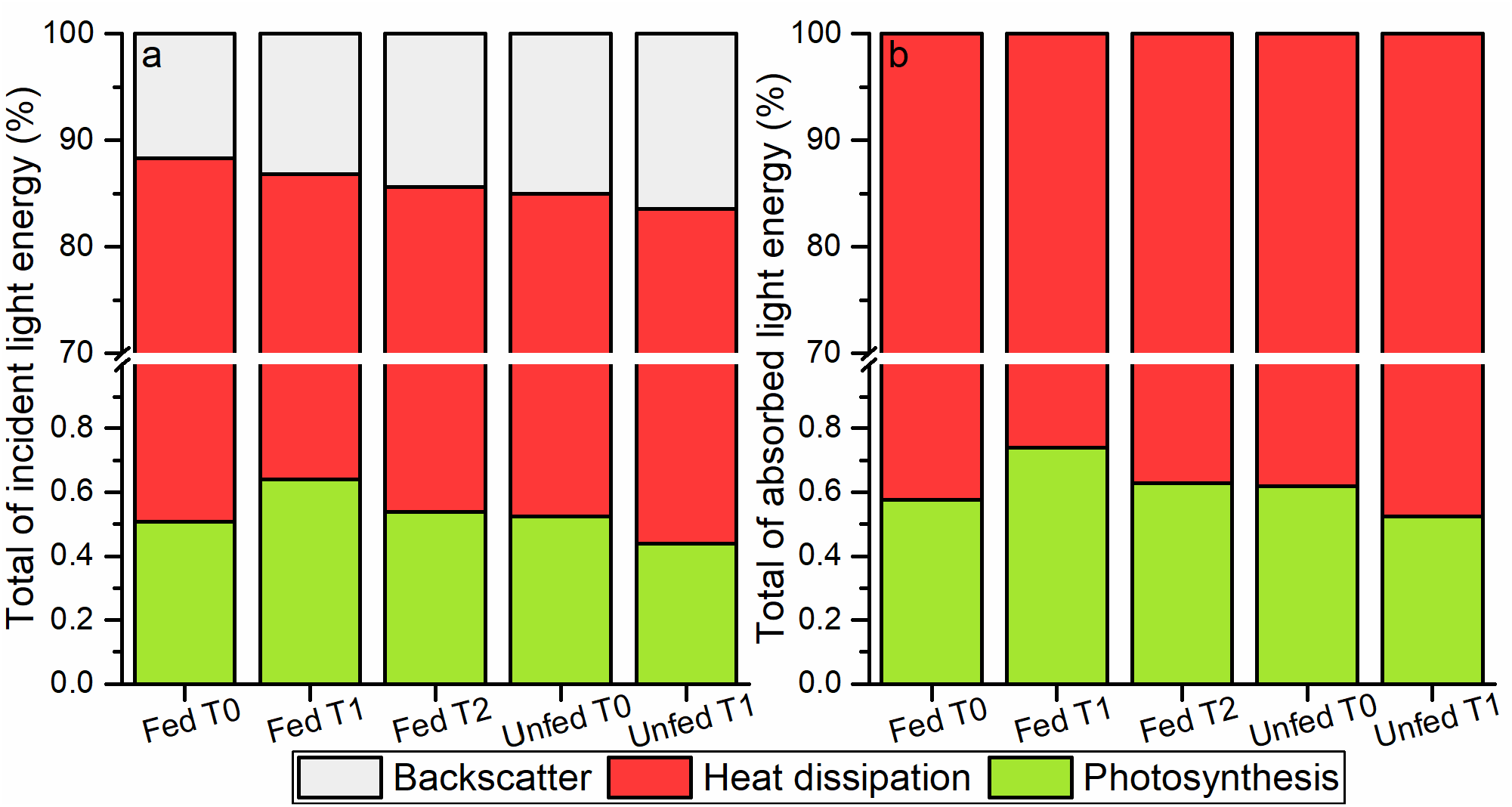
Radiative energy budget of fed and unfed *Pocillopora damicornis* under thermal stress in % of total incident (a) and total absorbed (b) light energy under high downwelling photon irradiance of 2400 μmol photons m^-2^ s^-1^. Gray bars indicate the amount of reflected light energy, red bars indicate the amount of light energy dissipated as heat, and green bars indicate amount of light energy conserved by photosynthesis. Note break in y-axis.

We calculated a theoretical radiative energy budget for a range of incident photon irradiance levels (from 80 to 2400 μmol photons m^-2^ s^-1^) based on detailed gross photosynthesis measurements on control fragments performed at these light levels (see methods). These extrapolated radiative energy budgets indicated that an increasing amount of light energy could be conserved by photosynthesis in both fed and unfed fragments, as the incident irradiance decreased and light-saturation of photosynthesis was alleviated (Figure 10). In fed fragments of *P. damicornis* the highest amount of photosynthetic energy use, i.e., about 5-6% of absorbed irradiance, was found at 80 μmol photons m^-2^ s^-1^, while unfed *P. damicornis* reached about 4% at T_0_. Corals from both treatments experienced an exponential decrease in photosynthetic energy quenching at higher irradiances, and reached the lowest amount of 0.5% for fed and 0.6% for unfed at 2400 μmol photons m^-2^ s^-1^. Photosynthetic use of absorbed light decreased to about 2.5% of absorbed light energy for unfed fragments after 3 days of thermal stress (T_1_), while fed fragments remained unaffected by thermal stress for 8 days (T_2_) before photosynthetic energy quenching dropped to about 4% (Figure 10).

**Figure 10.**
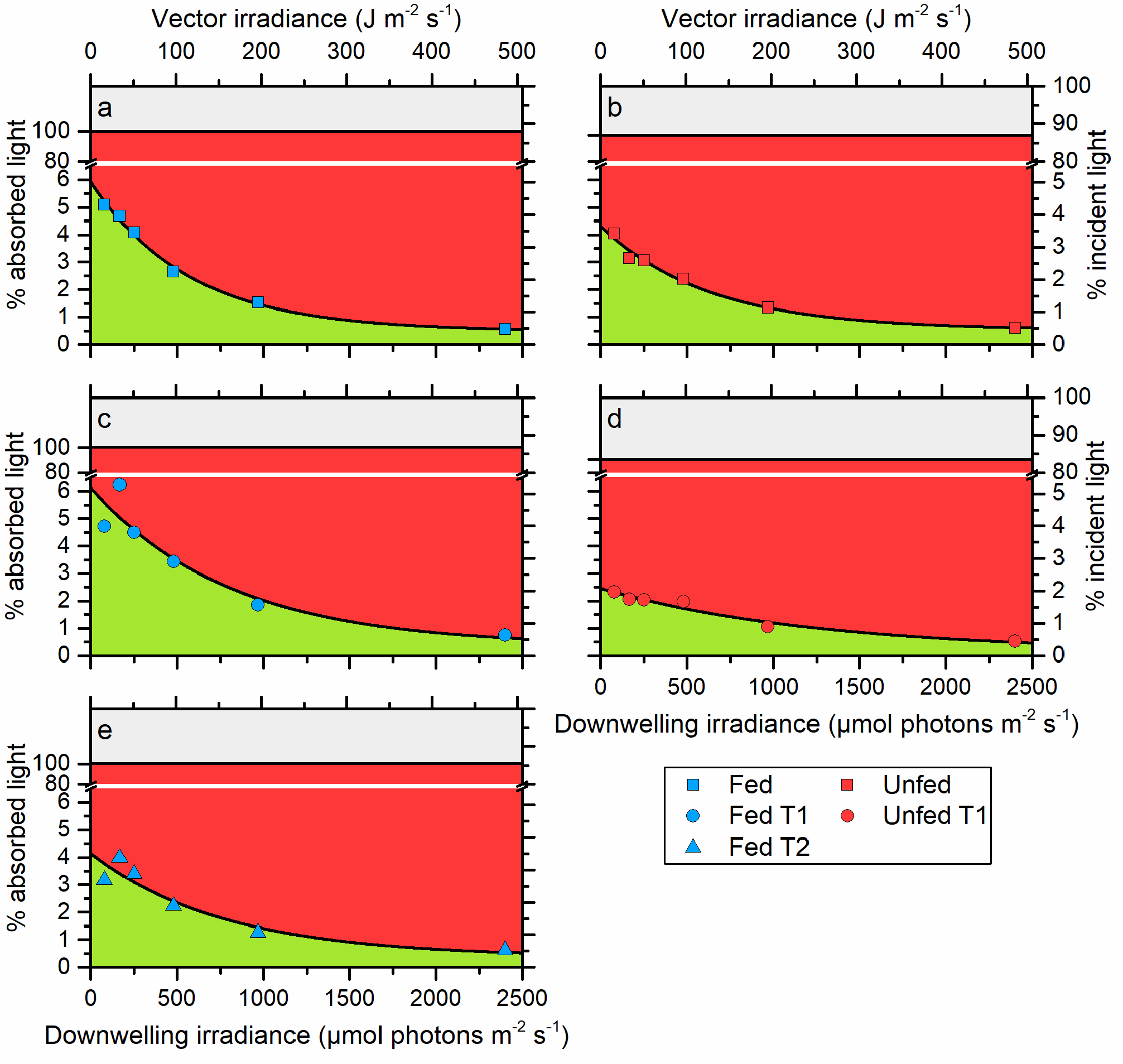
Calculated energy budgets in % of total absorbed light energy and in % of total incident light energy for fed (a, c, e) and unfed (b, d) *Pocillopora damicornis* without thermal stress (T_0_). Gray areas indicate the amount of reflected light energy, red areas indicate the amount of light energy dissipated as heat, and green areas indicate amount of light energy conserved by photosynthesis. Note break in y-axes.

## Discussion

In this study, we present the first directly linked measurements of the radiative energy and carbon assimilation budgets in the symbiont-bearing coral *Pocillopora damicornis*, and we compare the light energy utilization under thermal stress in fed and starved corals. We found that even though both fed and unfed corals responded to thermal stress by bleaching (Figure 1 and 5), the remaining zooxanthellae in both feeding treatments appeared photosynthetically competent (Figure 6b and 10).

### Radiative energy budgets

The balanced radiative energy budgets showed that fed and unfed corals absorb close to equal amounts of the incident light energy (85-88%; T_0_; Figure 9a). The amount of absorbed light energy decreased in both starved and fed corals upon thermal stress-induced bleaching. All radiative energy budgets showed highest energy dissipation as heat, where >99% of absorbed light energy was converted into heat regardless of feeding treatment and exposure to thermal stress, at the highest experimental irradiance of 2400 μmol photons m^-2^ s^-1^ (Figure 9b). The amount of absorbed light energy conserved by photosynthesis was very low (0.6-0.7%) under high irradiance (2400 μmol photons m^-2^ s^-1^) and remained virtually unchanged in corals from both treatments, regardless of exposure to thermal stress (Figure 9).

Since our study was conducted at a single, very high, photon irradiance (2400 μmol photons m^-2^ s^-1^) to obtain detectable TBLs for quantifying heat exchange, we estimated a theoretical radiative energy budget for 6 different lower levels of irradiance, where photosynthesis measurements were done on fragments from both treatments (See methods). Extrapolation of the radiative energy budget to lower incident irradiances showed that photosynthesis in control fragments accounted for up to ~4% of absorbed light energy in unfed corals and ~5-6% in fed corals under light-limited photosynthetic conditions (Figure 10a-b). This increased light use in fed corals suggests that fed *P. damicornis* had a higher photosynthetic efficiency when exposed to optimal temperature (25°C) and light conditions (<500 μmol photons m^-2^ s^-1^), compared to other coral species (e.g. Stylophora) (Ferrier-Pagès *et al*., 2010). Fed corals appeared unaffected by the temperature increase 3 days after the onset of thermal stress, whereas unfed corals expressed an immediate sign of stress via decreased photosynthetic quenching of absorbed light energy (4% to 2-2.5%; Figure 10d). Only after an additional 5 days of thermal stress did fed corals also express signs of stress by decreasing their photosynthetic quenching of absorbed light energy from 5-6% down to ~4% (Figure 10e).

The energy use efficiency of photosynthesis in highly pigmented photosynthetic biofilms are generally low due to their uniform topography and high optical density (Al-Najjar *et al*., 2010, 2012; Lichtenberg *et al*., 2017). In contrast, Brodersen *et al*. (2014) showed that the hemispherical coral *Montastrea curta* performed exceedingly better compared to other benthic photosystems, with a maximum photosynthetic energy efficiency of ~4% at 640 μmol photons m^-2^ s^-1^ under a flow velocity of 0.4 cm s^-1^. In our study, fed *P. damicornis* reach a photosynthetic energy efficiency of ~2% under a similar irradiance and a slightly slower flow velocity of 0.25 cm s^-1^. The difference in photosynthetic efficiency between *P. damicornis* and *M. curta* might be due to differences in skeletal heat conduction (Jimenez *et al*., 2012), as thin-tissued branching corals such as *P. damicornis* exhibit a higher heat conduction into the skeleton than thick-tissued massive corals.

### Photosynthesis and thermal stress

*Symbiodinium* density was steadily decreasing with thermal stress (Figure 3a), but chlorophyll content per cell and areal photosynthesis remained constant in fed and unfed corals (Figure 3c and 6a). High *Symbiodinium* cell densities (>1 x 10^6^ cells cm^-2^) can lead to algal self-shading (Enríquez *et al*., 2005), and thus reduced photosynthesis in deeper tissue layers due to high light attenuation. Loss of zooxanthellae during coral bleaching can initially alleviate such self-shading effects, leading to higher photosynthesis with fewer symbiont cells (Enríquez *et al*., 2005; Wangpraseurt *et al*., 2017b). This effect is also visible in our photosynthesis versus photon irradiance curves, where P_max_ remained constant (~0.50-0.52 nmol O_2_ cm^-2^ s^-1^) between thermal treatments (T_2_ relative to T_0_), while the α-value was considerably decreased after thermal stress (Figure 7a). This indicates that fed corals can uphold photosynthesis when thermally bleached, but they need higher irradiances to do so.

When comparing across all time points, the areal gross photosynthesis in fed corals was about 1.2 times higher than in unfed corals (Figure 6a). Feeding of corals has previously been shown to alleviate photodamage of *Symbiodinium*, where starved corals showed a decline in their nocturnal recovery rates of PSII relative to fed corals, and thus suffered more from chronic photoinhibition (Borell & Bischof, 2008; Borell *et al*., 2008). Unfed corals experienced a decrease in photosynthetic electron transport rate and net photosynthesis in addition to bleaching when thermally stressed relative to fed corals (Ferrier-Pagès *et al*., 2010). We found a similar pattern between our feeding treatments, as both relative electron transport rates and the effective yield of PSII was higher in fed relative to unfed corals before and after thermal stress (Figure).

We found that both rETR and Y(II) in fed corals increased with thermal stress (T_2_; Figure) which stands in contrast to the results from previous studies (e.g. Ferrier-Pagès *et al*., 2010). Thermal stress is known to inhibit the photosynthetic capabilities of aquatic phototrophs, as increased temperature leads to degradation of enzymes crucial for sustaining electron transport in the photosystems (Falkowski & Raven, 2007). However, our corals showed little to no decline in photosynthetic rates when exposed to thermal stress, and cell-specific photosynthesis rates of *Symbiodinium* increased by about a factor of 1.1 in unfed corals, and 1.5 in fed corals, after thermal stress (T_1_-T_2_ relative to T_0_; Figure 6b).

Comparisons across all treatments showed that cell specific gross photosynthesis was 2-fold higher in unfed corals relative to fed corals in the control group (T_0_), and almost 1.5 times higher after thermal stress (T_1_; Figure 6b). Even though coral cell density was based on the entire tissue area of the individual fragment (Figure 3), measurements were not performed entirely random (see methods), thus introducing a small error by apparently enhancing cell-specific gross photosynthesis in unfed corals due to a more heterogeneous distribution of symbionts in unfed coral fragment tissue (Figure 1). This issue also becomes apparent when comparing the high spatial resolution microsensor-based photosynthesis measurements from a single polyp, to optode measurements performed on entire coral fragments (See figure 2 in accompanying paper by Lyndby et al., submitted).

### Temperature microenvironment

Temperature microsensors measured a somewhat similar TBL thickness for unfed (~0.616 μm ±37.5) and fed corals (~0.745 μm ±67; Figure 8a and Supplementary figure 3). In contrast, the TBL of the massive thick-tissues coral *Montastrea curta* is >3 mm thick under comparable flow conditions (about 0.4 cm s^-1^; Brodersen et al., 2014). TBL thickness is controlled by coral growth form and microtopography (Jimenez *et al*., 2008). The small-polyped *P. damicornis* has a rather smooth tissue surface structure compared to massive faviid corals (Wangpraseurt *et al*., 2017a), thus explaining the relatively thin TBL’s found in *P. damicornis* (Jimenez *et al*., 2011).

Corals from either feeding treatment did not express any significant changes in radiative surface tissue warming (ΔT) as a result from thermal stress (Figure 8b). However, variable chlorophyll fluorescence imaging data of fed corals did reveal a clear decrease in NPQ after 8 days at 30°C (T_2_) relative to the control corals (T_0_, Supplementary figure 2). Spectral reflectance was increased for bleached corals, thus reducing the amount of absorbed light energy and coral surface warming (Enríquez *et al*., 2005; Jimenez *et al*., 2012). We found the same trend of spectral reflectance steadily increasing with thermal stress in corals from both feeding treatments (Figure 5), along with a clear negative correlation between coral spectral reflectance and symbiont density (R^2^ = 0.99; Supplementary figure 4). As such, the decrease in surface tissue heat exchange can be explained by the observed steady decrease in symbiont density and chlorophyll content during thermal stress (Figure 3).

### Light microenvironment

Scalar irradiance at the coral tissue surface was enhanced reaching ~120-130% of the incident downwelling irradiance for control corals (T_0_) in both feeding treatments. The scalar irradiance showed enhancements of up to ~140-145% of incident downwelling irradiance at the coral tissue surface after 3 days of thermal stress (T_1_; Figure 4a). Such enhancement of scalar irradiance in the coral tissue surface is common in bleached corals, where enhancements of PAR of up to 200% have previously been measured in polyp surface tissue of bleached *P. damicornis* (Wangpraseurt *et al*., 2017b).

The thin tissue of *P. damicornis* did not exhibit strong light gradients compared to thick-tissued corals (Wangpraseurt *et al*., 2012), and the scalar irradiance at the tissue-skeleton interface in polyp corallites was slightly lower compared to surface layers (95% of incident irradiance in fed, and 110% in unfed, T_0_, Figure 4b). This attenuation became even more apparent after the onset of thermal bleaching, as scalar irradiance at the tissue-skeleton interface in corals from both treatments further decreased (100% in unfed T_1_, and 78% in fed T_2_; Figure 4b). The reduced light attenuation during bleaching could be caused by the presence of endolithic microbes residing in the coral skeleton, where endolithic algal light absorption would decrease skeleton backscatter in certain regions of the light spectrum (Figure 4; compare T_1_ in panel c+f, and T_2_ in panel b+d; Fork & Larkum, 1989; Magnusson *et al*., 2007). However, no such endoliths appeared present to the naked eye.

## Conclusion

Based on our radiative energy budgets measured on fed and unfed corals during thermal stress-induced bleaching, we conclude that actively feeding *P. damicornis* can handle thermal stress better and apparently can maintain their energy demand as remaining symbionts show increased photosynthetic rates during bleaching. How such dynamics in symbiont numbers and photosynthesis affect the carbon budget in fed and starved *P. damicornis* corals during bleaching is investigated in an accompanying study (Lyndby et al., submitted).

## Acknowledgements

We thank the staff at the Scientific Center of Monaco for their excellent assistance with experimental procedures, and labor intensive general maintenance and care of the corals for the duration of this study. The study was funded by grants from the Carlsberg Foundation (DW, MK), a Sapere Aude Advanced Grant from the Independent Research Fund Denmark (DFF) (MK), and the Scientific Center of Monaco (part of the RTPI Nutress).

